# Interactive data analysis and reporting with SCIFR: A Single-File SPA Approach

**DOI:** 10.1101/2025.07.10.664259

**Authors:** Budi Permana, Thom P. Cuddihy, Brian M. Forde

## Abstract

The increasing complexity and scale of genomics data demand reporting solutions that are both interactive and easy to share. Here, we present SCIFR, a framework for creating self-contained, single-file interactive reports. SCIFR merges template-based flexibility with web app interactivity. Demonstrated with BLITSFR (comparative genomics) and METAXSFR (metagenomics), SCIFR outperforms established frameworks in efficiency and usability. The platform and tools are openly available, enabling next-generation, shareable reporting across diverse applications.

## Main

Increasing volumes and complexity of genomic data present significant challenges in interpreting results and communicating findings [1, 2]. Bioinformatics tools and pipelines increasingly feature interactive reports to enhance data exploration and interpretation. For example, MultiQC [3], Krona [4] and Clinker [5] compile results from analyses into interactive files, enabling users to efficiently review and interrogate diverse datasets. These interactive capabilities provide a richer, more flexible environment for data interpretation, surpassing the limitations of static tables or figures that offer a limited view of the data.

Common strategies for bioinformatics interactive reporting typically fall into two categories: static report generation and dynamic web applications. Static approaches, exemplified by MultiQC and Clinker [5, 6], use template engines such as Jinja to process data through predefined templates and generate HTML files. Web applications, such as Proksee and Shiny-based applications [7-10], employ server-client architecture to dynamically generate content in response to user interactions. While the former integrate seamlessly with command-line workflows due to their self-contained nature, the latter offer enhanced functionality but require a web server that often complicates integration with typical HPC-based bioinformatics pipelines. However, recent advances in single-page application (SPA) frameworks have enabled web developers to create sophisticated and efficient client-side web applications [11-13]. Unlike traditional web applications, which require server infrastructure, these SPAs bundle their logic, styling, and data into distributable packages that run entirely in the browser.

Here, we introduce SCIFR, a development framework that leverages SPA to generate self-contained, interactive single-file reports. SCIFR outputs a single HTML file with advanced features including client-side routing, state management processing, and dynamic visualisation, hence featuring both templating and web application capabilities without requiring a web server. By combining these features and leveraging the growing ecosystem of bioinformatics-focused JavaScript libraries [7, 14-16], SCIFR enables versatile, interactive reporting for a wide range of bioinformatics applications.

In the SCIFR system, the SPA build acts as the template *Viewer* (Figure 1b), responsible for rendering outputs from the *Processor* (Figure 1c). This *Processor* can range from a simple script to a complex pipeline generating text data for presentation in the *Viewer*. Integration between the *Processor* and the *Viewer* is managed by the *Mutator* (Figure 1d), which identifies uniquely tagged input placeholders within the template and replaces them with new data to produce the final HTML report (Figure 1e). To support this system workflow, we provide a reusable boilerplate built using React.js and additional JavaScript libraries as the template backbone (Figure 1a, Supplementary table S1). The boilerplate enables rapid deployment of the development environment and is freely available via the Node.js package manager. The *Mutator* and example *Processor* pipelines are also accessible in the GitHub repository (https://github.com/nalarbp/scifr).

**Figure 1.**
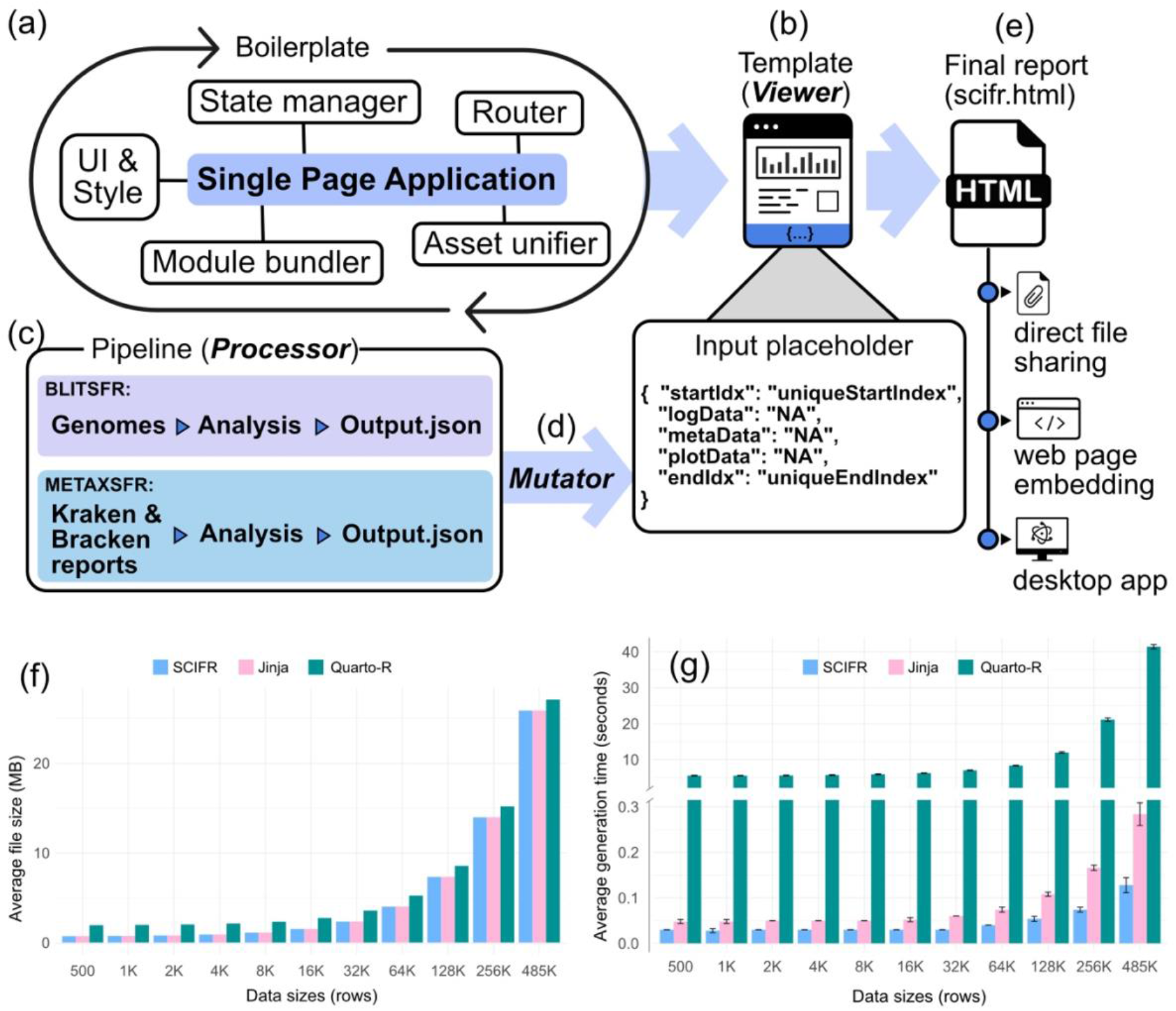
Overview of SCIFR components and workflow. (a) SCIFR boilerplate allows developers to develop SCIFR templates through rapid setup of single page application environments and components. The resulting template, which serves as the *Viewer*, contains placeholder JSON input text that can be replaced to produce new reports (b). Example pipelines presented in this study (BLITSFR and METAXSFR) function as the central *Processors* (c), producing data to be transferred by the *Mutator* (d) to the template, resulting in a final HTML report (e). The average report file size in megabytes (f) and report generation time, in seconds, (g) when benchmarked against Jinja and Quarto-R workflows.

To evaluate SCIFR’s performance within the existing report generation ecosystem, we benchmarked it against two established workflows: Jinja (Python-based; https://github.com/pallets/jinja) and Quarto (R-version; https://github.com/quarto-dev/quarto-cli). We generated interactive reports featuring identical charts and tables visualising COVID-19 surveillance data [17] across varying dataset sizes (Supplementary figure S1). For each workflow and dataset, we measured report generation time and output file size (Supplementary table S2).

All workflows generated files of comparable size, which scaled linearly with the volume of input data (Figure 1f), highlighting that data input embedding is the primary determinant of the final report’s file size. However, SCIFR demonstrated exceptional speed in report generation, achieving an average performance 200 times faster than Quarto-R and closely matching Jinja (∼1.8 times faster) across all datasets (Figure 1g). This efficiency is attributed to SCIFR’s architecture, which separates template compilation from data injection. As a result, report generation primarily involves inserting data into pre-built templates. Furthermore, SCIFR’s JavaScript-based workflow removes the translation overhead in Quarto-R and Jinja when converting data structures from their native environments into JavaScript code for browser compatibility.

Next, we demonstrated SCIFR’s applicability by developing two bioinformatics tools: BLITSFR (BLAST Interactive Tracks in Single File Report) and METAXSFR (Metagenome Taxonomic Explorer in Single File Report).

BLITSFR is a Nextflow-based command-line tool that compares query sequences against a reference sequence and visualises results via interactive tracks rendered using CGView.js [7, 18]. Its workflow includes designing a SCIFR template that compatible with CGview.js, running a Nextflow pipeline to align sequence queries to a reference using BLASTn [19] (for assemblies) or KMA [20] (for reads), and inserting the results into the template. The resulting BLITSFR report displays BLAST hits or read coverage plots as interactive tracks with options for metadata-driven colouring, filtering and sorting. To validate its functionality, we reanalysed sequence data from a previously described hospital outbreak of vancomycin-resistant *Enterococcus faecium* [21]. BLITSFR enabled rapid comparison, sorting, and colour-coding of query genomes according to lineage metadata, which facilitated swift identification of conserved and variable regions among sub-lineages (Figure 2a). The complete analysis of the dataset was performed in under 2 minutes on a standard notebook, and took less than 1 minute when executed on a HPC.

**Figure 2.**
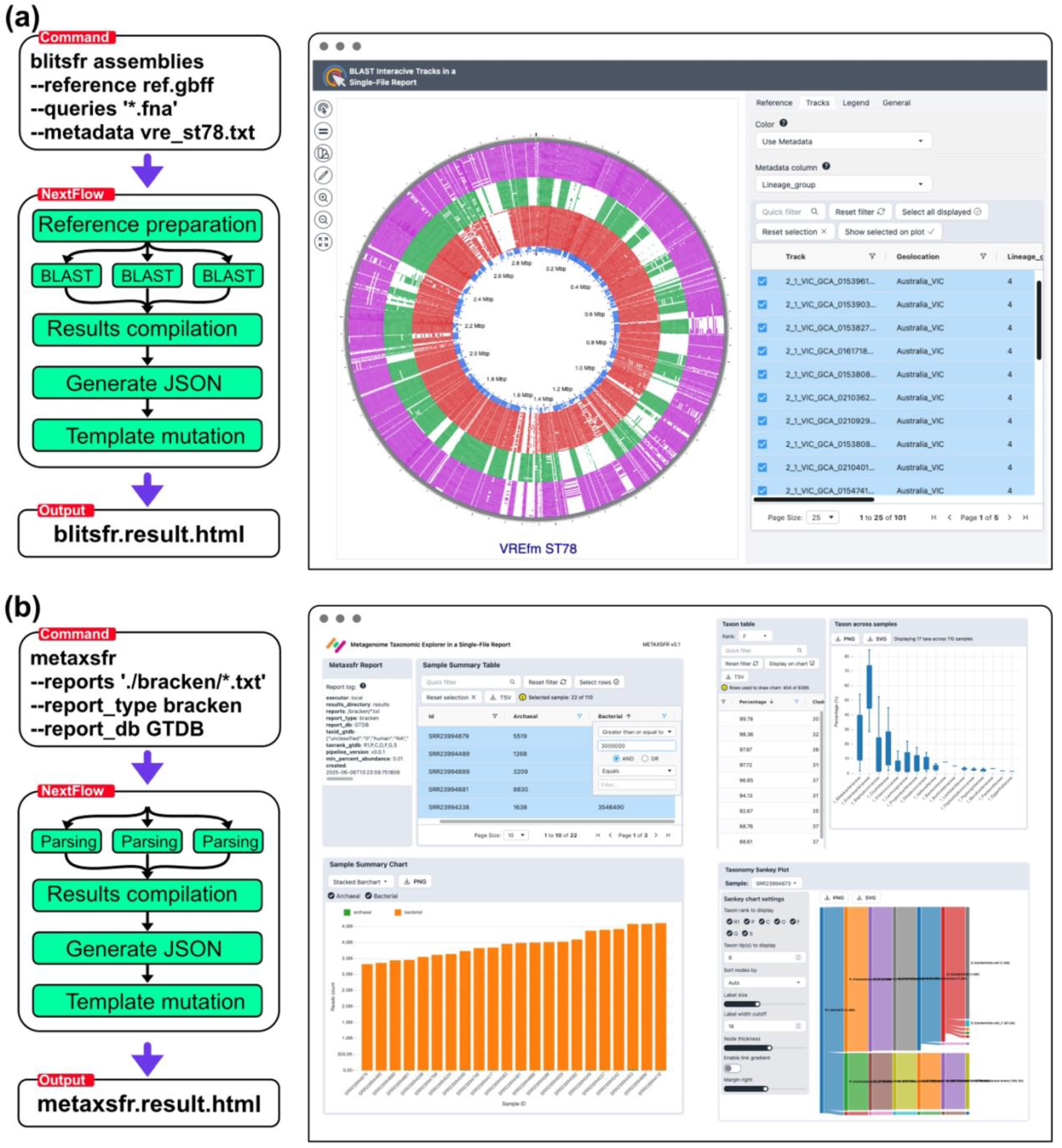
BLITSFR and METAXSFR pipelines using SCIFR to generate an interactive report. (a) BLITSFR report showing BLAST hits of 100 VREfm ST78 genomes [21] against a reference genome (SAMN39986858). The tracks are coloured and sorted based on genome lineage groups provided in the metadata. (b) METAXSFR report showing taxonomic profiles of 110 metagenome samples from Heston *et al*. [28].

METAXSFR, our second SCIFR implementation, aggregates and visualises taxonomic profiles generated by Kraken2 [22], Bracken [23] or Metaphlan4 [24]. Leveraging Nextflow for efficient large-scale data processing, it summarises and fills SCIFR template featuring interactive tables, charts and Sankey diagrams for exploring taxa abundance and taxonomic hierarchies, similar to Krona [4] and Pavian [25]. METAXSFR also supports both NCBI [26] and GTDB [27] taxonomic identifiers, ensuring broad compatibility. Its performance was validated by processing 110 Bracken reports from infant gut metagenome samples [27], which enabled a detailed exploration of community composition, in line with published findings [28]. Using this dataset, METAXSR produced an interactive HTML report in under 30 seconds, streamlining data investigation for users.

SCIFR opens new possibilities for bioinformatics tool development that were previously challenging or impractical. For instance, BLITSFR overcomes key limitations of existing comparative genomic visualisation tools such as BRIG [29], which only generates static images and requires time-consuming manual configuration before analysis, and Proksee [7] or CCT [18], which are centralised web services that necessitate uploading data to remote servers and prevent analysis on local resources. Similarly, METAXSFR addresses the automation limitations of Pavian, a Shiny application that cannot easily generate reports without user upload and interaction [25]. Additionally, SCIFR builds on established report generation strategies like MultiQC, which use templating and JavaScript to aggregate and visualise results [3], but leveraging SPA architecture for more sophisticated and dynamic visualisations. For example, BLITSFR enables real-time filtering and dynamic visualisation updates on BLAST results through advanced JavaScript applications state management.

As a self-contained single file report, SCIFR has several limitations. Firstly, the size of the report scales with the amount of input data (Figure 1f), which can be addressed by compressing the file or hosting large reports on a local network. Secondly, because all visualisation logic runs client-side, larger or complex datasets may exceed browser memory limit. To mitigate this, data can be pre-processed to reduce computational complexity and data pagination can be implemented to load subsets on demand.

In summary, we present SCIFR, a single-file interactive report development framework that facilitates data-analysis and creation of sophisticated, web application-like reports without server dependencies. SCIFR integrates seamlessly with bioinformatics pipelines and demonstrates substantial efficiency gains—generating reports up to 200 times faster than established frameworks. As bioinformatics datasets continue to grow in complexity, SCIFR provides a robust foundation for next-generation interactive reporting, supporting enhanced data interpretation and scientific collaboration.

## Methods

### Code and data availability

The SCIFR platform is showcased in https://scifr.fordelab.com. The boilerplate create-scifr is available in NPM repository (https://www.npmjs.com/package/create-scifr). BLITSFR and METAXSFR, along with test datasets, are available on GitHub (https://github.com/nalarbp/blitsfr; https://github.com/nalarbp/metaxsfr).

### Developing SCIR template

To initiate the development environment, the workflow uses create-react-app (https://github.com/facebook/create-react-app) followed by the installation of core JavaScript libraries, including Jotai for state management (https://github.com/pmndrs/jotai), TailwindCSS for user interface components and styling (https://github.com/tailwindlabs/tailwindcss), and HTML-inline Webpack plugins for asset unification (https://github.com/jantimon/html-webpack-plugin). A boilerplate for this environment setup has been created and is available at the public npm registry (https://www.npmjs.com/package/create-scifr). To use the boilerplate users can do: (1) *npx create-scif my_scifr_template* that will initiate development environment, download, and install required dependencies. (2) *npm run dev* to start the single-page-application (SPA) development server, and let developer create and change components to make a desired SPA. (3) *npm run build* will finally bundle the SPA into a single html template file.

### Benchmarking SCIFR

We developed functionally identical interactive reports using three HTML generation approaches: SCIFR (React-based), Jinja2 (Python templating), and Quarto-R (updated R-markdown approach). Each report implemented identical interactive features including a sortable data table using DataTables.js v1.13.4 and interactive charts using Chart.js v4.2.1, with all JavaScript libraries loaded locally to ensure consistency across tools. Reports were generated using a COVID-19 dataset [25] containing 485K original data points, systematically scaled to different sizes: 500, 1K, 2K, 4K, 8K, 16K, 32K, 64K, 128K, 256K, and 485K. Report generation time and output file size were measured and averaged across five iterations to account for system variability. All benchmarks were conducted on a MacBook Pro M3 with 16GB RAM, and a high performance computing cluster (HPC) with 24 CPUs and 8Gb RAM under controlled conditions, with system caches cleared and a delay applied between measurements sessions to ensure consistent baseline performance.

## Supplementary figure and tables

**Supplementary figure S1.**
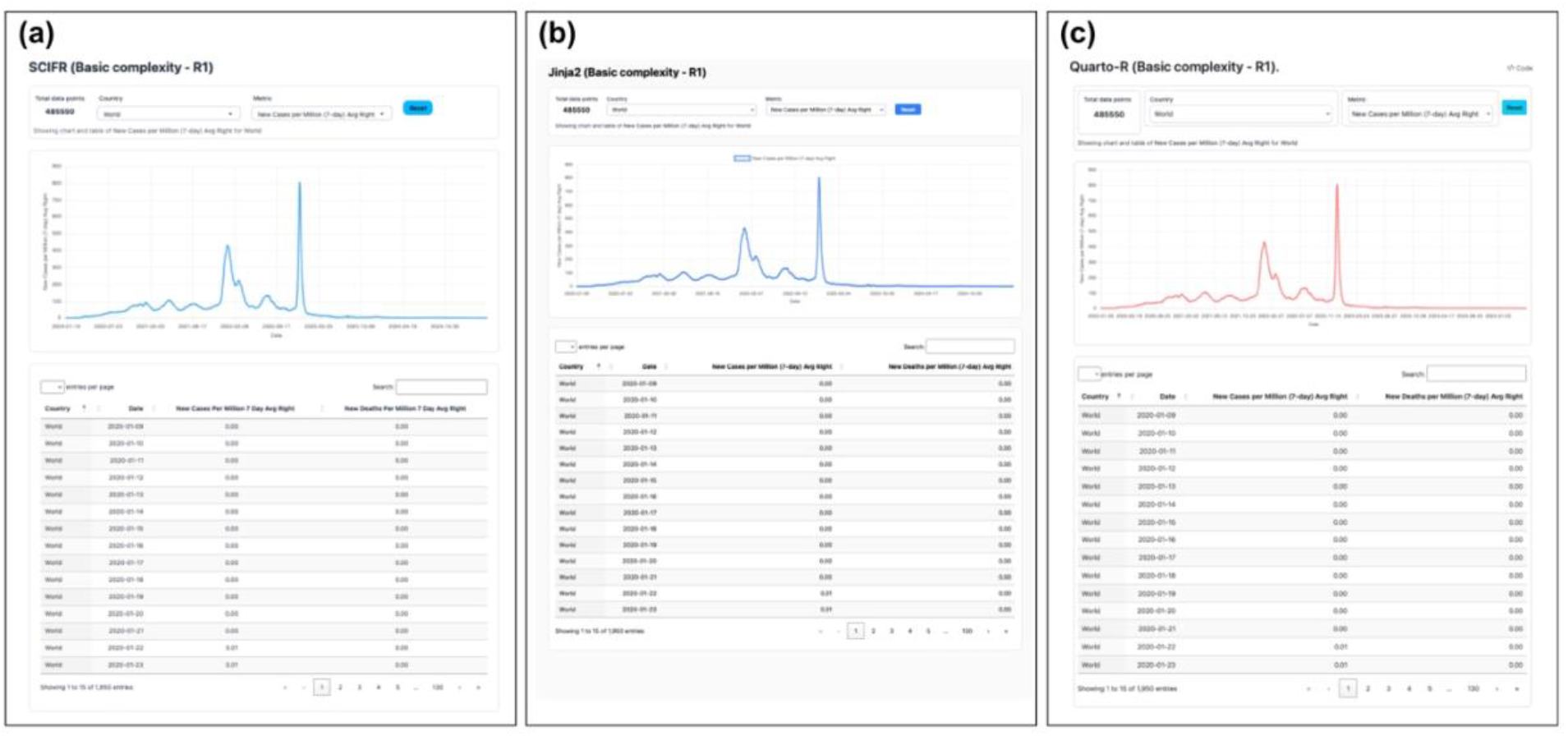
A functionally identical single file HTML report showing interactive chart and table of 485,550 COVID-19 cases generated using (a) SCIFR, (b) Jinja, and (c) Quarto-R. All interactive reports are available in https://github.com/nalarbp/scifr GitHub repository.

**Supplementary table S1.**
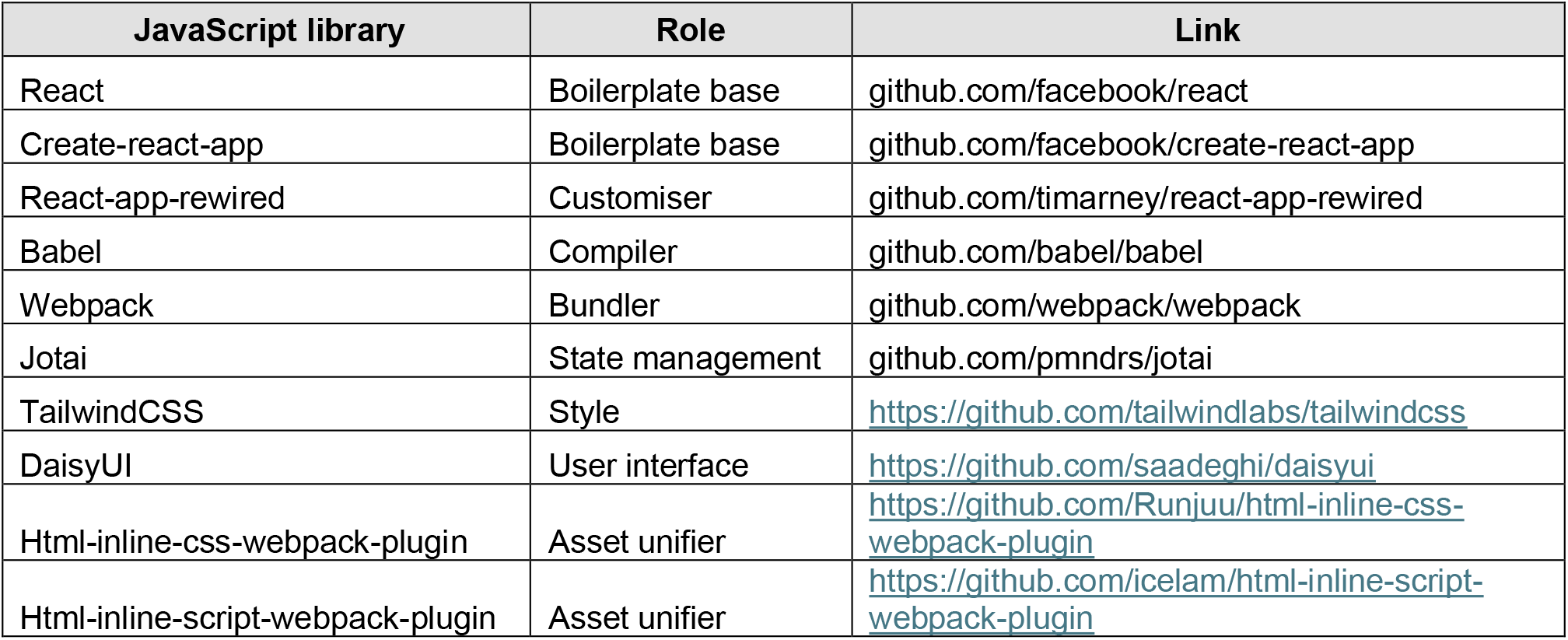
Core JavaScript libraries of SCIFR’s boilerplate.

**Supplementary table S2.** Report generation time and file size of a test single file HTML produced by SCIFR, Jinja (Python-based), and Quarto-R (R-markdown based).

| Tool | Test iteration | Data size (rows) | Generation time (seconds) | File size (bytes) |
| --- | --- | --- | --- | --- |
| quartoR | 1 | 500 | 5.57 | 1945176 |
| quartoR | 1 | 1K | 5.59 | 1971090 |
| quartoR | 1 | 2K | 5.54 | 2022925 |
| quartoR | 1 | 4K | 5.88 | 2126596 |
| quartoR | 1 | 8K | 5.91 | 2333967 |
| quartoR | 1 | 16K | 6.3 | 2748644 |
| quartoR | 1 | 32K | 7.03 | 3578155 |
| quartoR | 1 | 64K | 8.41 | 5236991 |
| quartoR | 1 | 128K | 12.22 | 8554689 |
| quartoR | 1 | 256K | 21.73 | 15190190 |
| quartoR | 1 | 485K | 42.05 | 27089856 |
| quartoR | 2 | 500 | 5.54 | 1945176 |
| quartoR | 2 | 1K | 5.49 | 1971090 |
| quartoR | 2 | 2K | 5.58 | 2022925 |
| quartoR | 2 | 4K | 5.59 | 2126596 |
| quartoR | 2 | 8K | 5.85 | 2333967 |
| quartoR | 2 | 16K | 6.2 | 2748644 |
| quartoR | 2 | 32K | 6.89 | 3578155 |
| quartoR | 2 | 64K | 8.31 | 5236991 |
| quartoR | 2 | 128K | 12.01 | 8554689 |
| quartoR | 2 | 256K | 21.42 | 15190190 |
| quartoR | 2 | 485K | 41.72 | 27089856 |
| quartoR | 3 | 500 | 5.5 | 1945176 |
| quartoR | 3 | 1K | 5.53 | 1971090 |
| quartoR | 3 | 2K | 5.76 | 2022925 |
| quartoR | 3 | 4K | 5.65 | 2126596 |
| quartoR | 3 | 8K | 5.86 | 2333967 |
| quartoR | 3 | 16K | 6.24 | 2748644 |
| quartoR | 3 | 32K | 7.1 | 3578155 |
| quartoR | 3 | 64K | 8.42 | 5236991 |
| quartoR | 3 | 128K | 11.85 | 8554689 |
| quartoR | 3 | 256K | 20.78 | 15190190 |
| quartoR | 3 | 485K | 40.75 | 27089856 |
| quartoR | 4 | 500 | 5.71 | 1945176 |
| quartoR | 4 | 1K | 5.64 | 1971090 |
| quartoR | 4 | 2K | 5.58 | 2022925 |
| quartoR | 4 | 4K | 5.72 | 2126596 |
| quartoR | 4 | 8K | 6.13 | 2333967 |
| quartoR | 4 | 16K | 6.3 | 2748644 |
| quartoR | 4 | 32K | 7.04 | 3578155 |
| quartoR | 4 | 64K | 8.28 | 5236991 |
| quartoR | 4 | 128K | 12.07 | 8554689 |
| quartoR | 4 | 256K | 20.86 | 15190190 |
| quartoR | 4 | 485K | 41.73 | 27089856 |
| quartoR | 5 | 500 | 5.42 | 1945176 |
| quartoR | 5 | 1K | 5.49 | 1971090 |
| quartoR | 5 | 2K | 5.49 | 2022925 |
| quartoR | 5 | 4K | 5.66 | 2126596 |
| quartoR | 5 | 8K | 5.75 | 2333967 |
| quartoR | 5 | 16K | 6.14 | 2748644 |
| quartoR | 5 | 32K | 6.89 | 3578155 |
| quartoR | 5 | 64K | 8.23 | 5236991 |
| quartoR | 5 | 128K | 11.66 | 8554689 |
| quartoR | 5 | 256K | 20.76 | 15190190 |
| quartoR | 5 | 485K | 40.86 | 27089856 |
| jinja2 | 1 | 500 | 0.04 | 723059 |
| jinja2 | 1 | 1K | 0.04 | 748987 |
| jinja2 | 1 | 2K | 0.05 | 800850 |
| jinja2 | 1 | 4K | 0.05 | 904577 |
| jinja2 | 1 | 8K | 0.05 | 1112060 |
| jinja2 | 1 | 16K | 0.06 | 1526954 |
| jinja2 | 1 | 32K | 0.06 | 2356913 |
| jinja2 | 1 | 64K | 0.07 | 4016652 |
| jinja2 | 1 | 128K | 0.1 | 7336149 |
| jinja2 | 1 | 256K | 0.16 | 13975248 |
| jinja2 | 1 | 485K | 0.26 | 25881368 |
| jinja2 | 2 | 500 | 0.05 | 723059 |
| jinja2 | 2 | 1K | 0.05 | 748987 |
| jinja2 | 2 | 2K | 0.05 | 800850 |
| jinja2 | 2 | 4K | 0.05 | 904577 |
| jinja2 | 2 | 8K | 0.05 | 1112060 |
| jinja2 | 2 | 16K | 0.05 | 1526954 |
| jinja2 | 2 | 32K | 0.06 | 2356913 |
| jinja2 | 2 | 64K | 0.07 | 4016652 |
| jinja2 | 2 | 128K | 0.11 | 7336149 |
| jinja2 | 2 | 256K | 0.16 | 13975248 |
| jinja2 | 2 | 485K | 0.27 | 25881368 |
| jinja2 | 3 | 500 | 0.05 | 723059 |
| jinja2 | 3 | 1K | 0.05 | 748987 |
| jinja2 | 3 | 2K | 0.05 | 800850 |
| jinja2 | 3 | 4K | 0.05 | 904577 |
| jinja2 | 3 | 8K | 0.05 | 1112060 |
| jinja2 | 3 | 16K | 0.05 | 1526954 |
| jinja2 | 3 | 32K | 0.06 | 2356913 |
| jinja2 | 3 | 64K | 0.08 | 4016652 |
| jinja2 | 3 | 128K | 0.11 | 7336149 |
| jinja2 | 3 | 256K | 0.17 | 13975248 |
| jinja2 | 3 | 485K | 0.27 | 25881368 |
| jinja2 | 4 | 500 | 0.05 | 723059 |
| jinja2 | 4 | 1K | 0.05 | 748987 |
| jinja2 | 4 | 2K | 0.05 | 800850 |
| jinja2 | 4 | 4K | 0.05 | 904577 |
| jinja2 | 4 | 8K | 0.05 | 1112060 |
| jinja2 | 4 | 16K | 0.05 | 1526954 |
| jinja2 | 4 | 32K | 0.06 | 2356913 |
| jinja2 | 4 | 64K | 0.07 | 4016652 |
| jinja2 | 4 | 128K | 0.11 | 7336149 |
| jinja2 | 4 | 256K | 0.17 | 13975248 |
| jinja2 | 4 | 485K | 0.3 | 25881368 |
| jinja2 | 5 | 500 | 0.05 | 723059 |
| jinja2 | 5 | 1K | 0.05 | 748987 |
| jinja2 | 5 | 2K | 0.05 | 800850 |
| jinja2 | 5 | 4K | 0.05 | 904577 |
| jinja2 | 5 | 8K | 0.05 | 1112060 |
| jinja2 | 5 | 16K | 0.05 | 1526954 |
| jinja2 | 5 | 32K | 0.06 | 2356913 |
| jinja2 | 5 | 64K | 0.08 | 4016652 |
| jinja2 | 5 | 128K | 0.11 | 7336149 |
| jinja2 | 5 | 256K | 0.17 | 13975248 |
| jinja2 | 5 | 485K | 0.32 | 25881368 |
| scifr | 1 | 500 | 0.03 | 717586 |
| scifr | 1 | 1K | 0.02 | 743506 |
| scifr | 1 | 2K | 0.03 | 795353 |
| scifr | 1 | 4K | 0.03 | 899048 |
| scifr | 1 | 8K | 0.03 | 1106467 |
| scifr | 1 | 16K | 0.03 | 1521237 |
| scifr | 1 | 32K | 0.03 | 2350940 |
| scifr | 1 | 64K | 0.04 | 4010163 |
| scifr | 1 | 128K | 0.05 | 7328632 |
| scifr | 1 | 256K | 0.07 | 13965675 |
| scifr | 1 | 485K | 0.11 | 25868107 |
| scifr | 2 | 500 | 0.03 | 717586 |
| scifr | 2 | 1K | 0.03 | 743506 |
| scifr | 2 | 2K | 0.03 | 795353 |
| scifr | 2 | 4K | 0.03 | 899048 |
| scifr | 2 | 8K | 0.03 | 1106467 |
| scifr | 2 | 16K | 0.03 | 1521237 |
| scifr | 2 | 32K | 0.03 | 2350940 |
| scifr | 2 | 64K | 0.04 | 4010163 |
| scifr | 2 | 128K | 0.05 | 7328632 |
| scifr | 2 | 256K | 0.07 | 13965675 |
| scifr | 2 | 485K | 0.15 | 25868107 |
| scifr | 3 | 500 | 0.03 | 717586 |
| scifr | 3 | 1K | 0.03 | 743506 |
| scifr | 3 | 2K | 0.03 | 795353 |
| scifr | 3 | 4K | 0.03 | 899048 |
| scifr | 3 | 8K | 0.03 | 1106467 |
| scifr | 3 | 16K | 0.03 | 1521237 |
| scifr | 3 | 32K | 0.03 | 2350940 |
| scifr | 3 | 64K | 0.04 | 4010163 |
| scifr | 3 | 128K | 0.06 | 7328632 |
| scifr | 3 | 256K | 0.08 | 13965675 |
| scifr | 3 | 485K | 0.12 | 25868107 |
| scifr | 4 | 500 | 0.03 | 717586 |
| scifr | 4 | 1K | 0.03 | 743506 |
| scifr | 4 | 2K | 0.03 | 795353 |
| scifr | 4 | 4K | 0.03 | 899048 |
| scifr | 4 | 8K | 0.03 | 1106467 |
| scifr | 4 | 16K | 0.03 | 1521237 |
| scifr | 4 | 32K | 0.03 | 2350940 |
| scifr | 4 | 64K | 0.04 | 4010163 |
| scifr | 4 | 128K | 0.06 | 7328632 |
| scifr | 4 | 256K | 0.08 | 13965675 |
| scifr | 4 | 485K | 0.14 | 25868107 |
| scifr | 5 | 500 | 0.03 | 717586 |
| scifr | 5 | 1K | 0.03 | 743506 |
| scifr | 5 | 2K | 0.03 | 795353 |
| scifr | 5 | 4K | 0.03 | 899048 |
| scifr | 5 | 8K | 0.03 | 1106467 |
| scifr | 5 | 16K | 0.03 | 1521237 |
| scifr | 5 | 32K | 0.03 | 2350940 |
| scifr | 5 | 64K | 0.04 | 4010163 |
| scifr | 5 | 128K | 0.05 | 7328632 |
| scifr | 5 | 256K | 0.07 | 13965675 |
| scifr | 5 | 485K | 0.12 | 25868107 |

## Abbreviations

BLAST: Basic Local Alignment Search Tool
BLITSFR: BLAST Interactive Tracks in Single File Report
GTDB: Genome Taxonomy Database
HPC: High-performance computing
HTML: Hyper Text Markup Language
METAXSFR: Metagenome Taxonomic Explorer in Single File Report
NCBI: National Center for Biotechnology Information
SCIFR: Self Contained Interactive Single-File Report
SPA: Single Page Application
RAM: Random Access Memory

## References

1. O’Donoghue, S.I, Grand Challenges in Bioinformatics Data Visualization. Front Bioinform, 2021. 1: p. 669186.

2. Tao, Y., et al., Information Visualization Techniques in Bioinformatics during the Postgenomic Era. Drug Discov Today Biosilico, 2004. 2(6): p. 237–245.

3. Ewels, P., et al., MultiQC: summarize analysis results for multiple tools and samples in a single report. Bioinformatics, 2016. 32(19): p. 3047–8.

4. Ondov, B.D., N.H. Bergman, and A.M. Phillippy, Interactive metagenomic visualization in a Web browser. BMC Bioinformatics, 2011. 12: p. 385.

5. Gilchrist, C.L.M. and Y.H. Chooi, clinker & clustermap.js: automatic generation of gene cluster comparison figures. Bioinformatics, 2021. 37(16): p. 2473–2475.

6. Rashid, U., et al., AssemblyQC: A Nextflow pipeline for reproducible reporting of assembly quality. Bioinformatics, 2024.

7. Grant, J.R., et al., Proksee: in-depth characterization and visualization of bacterial genomes. Nucleic Acids Res, 2023. 51(W1): p. W484–W492.

8. Argimon, S., et al., A global resource for genomic predictions of antimicrobial resistance and surveillance of Salmonella Typhi at pathogenwatch. Nat Commun, 2021. 12(1): p. 2879.

9. Argimon, S., et al., Microreact: visualizing and sharing data for genomic epidemiology and phylogeography. Microb Genom, 2016. 2(11): p. e000093.

10. Chang, W., et al., shiny: Web Application Framework for R. 2024.

11. Hadfield, J., et al., Phandango: an interactive viewer for bacterial population genomics. Bioinformatics, 2018. 34(2): p. 292–293.

12. Permana, B., S.A. Beatson, and B.M. Forde, GraphSNP: an interactive distance viewer for investigating outbreaks and transmission networks using a graph approach. BMC Bioinformatics, 2023. 24(1): p. 209.

13. Permana, B., et al., HAIviz: an interactive dashboard for visualising and integrating healthcare-associated genomic epidemiological data. Microb Genom, 2024. 10(2).

14. Robinson, J.T., et al., igv.js: an embeddable JavaScript implementation of the Integrative Genomics Viewer (IGV). Bioinformatics, 2023. 39(1).

15. Franz, M., et al., Cytoscape.js 2023 update: a graph theory library for visualization and analysis. Bioinformatics, 2023. 39(1).

16. Shank, S.D., S. Weaver, and S.L. Kosakovsky Pond, phylotree.js - a JavaScript library for application development and interactive data visualization in phylogenetics. BMC Bioinformatics, 2018. 19(1): p. 276.

17. Mathieu, E., et al., Coronavirus (COVID-19) Cases. Our World in Data, 2020.

18. Stothard, P., J.R. Grant, and G. Van Domselaar, Visualizing and comparing circular genomes using the CGView family of tools. Brief Bioinform, 2019. 20(4): p. 1576–1582.

19. Camacho, C., et al., BLAST+: architecture and applications. BMC Bioinformatics, 2009. 10: p. 421.

20. Clausen, P., F.M. Aarestrup, and O. Lund, Rapid and precise alignment of raw reads against redundant databases with KMA. BMC Bioinformatics, 2018. 19(1): p. 307.

21. Permana, B., et al., Using Genomics To Investigate an Outbreak of Vancomycin-Resistant Enterococcus faecium ST78 at a Large Tertiary Hospital in Queensland. Microbiol Spectr, 2023. 11(3): p. e0420422.

22. Wood, D.E., J. Lu, and B. Langmead, Improved metagenomic analysis with Kraken 2. Genome Biol, 2019. 20(1): p. 257.

23. Lu, J., et al., Bracken: estimating species abundance in metagenomics data. PeerJ Comput Sci, 2017. 3.

24. Blanco-Miguez, A., et al., Extending and improving metagenomic taxonomic profiling with uncharacterized species using MetaPhlAn 4. Nat Biotechnol, 2023. 41(11): p. 1633–1644.

25. Breitwieser, F.P. and S.L. Salzberg, Pavian: interactive analysis of metagenomics data for microbiome studies and pathogen identification. Bioinformatics, 2020. 36(4): p. 1303–1304.

26. Federhen, S., The NCBI Taxonomy database. Nucleic Acids Res, 2012. 40(Database issue): p. D136–43.

27. Parks, D.H., et al., GTDB: an ongoing census of bacterial and archaeal diversity through a phylogenetically consistent, rank normalized and complete genome-based taxonomy. Nucleic Acids Research, 2022. 50(D1): p. D785–D794.

28. Heston, S.M., et al., Strain-resolved metagenomic analysis of the gut as a reservoir for bloodstream infection pathogens among premature infants in Singapore. Gut Pathogens, 2023. 15(1).

29. Alikhan, N.F., et al., BLAST Ring Image Generator (BRIG): simple prokaryote genome comparisons. BMC Genomics, 2011. 12: p. 402.

